# Oncogenic signaling inhibits c-FLIP expression and promotes cancer cell survival during ECM-detachment

**DOI:** 10.1101/2021.04.19.440544

**Authors:** Matyas Abel Tsegaye, Jianping He, Kyle McGeehan, Ireland M. Murphy, Mati Nemera, Zachary T. Schafer

## Abstract

Inhibition of programmed cell death pathways is frequently observed in cancer cells where it functions to facilitate tumor progression. However, some proteins involved in the regulation of cell death function dichotomously to both promote and inhibit cell death depending on the cellular context. As such, understanding how cell death proteins are regulated in a context-dependent fashion in cancer cells is of utmost importance. We have uncovered evidence that cellular FLICE-like Inhibitory Protein (c-FLIP), a well-known anti-apoptotic protein, is often downregulated in tumor tissue when compared to adjacent normal tissue. These data argue that c-FLIP may have activity distinct from its canonical role in antagonizing cell death. Interestingly, we have discovered that detachment from extracellular matrix (ECM) serves as a signal to elevate c-FLIP transcription and that oncogenic signaling blocks ECM-detachment-induced c-FLIP elevation. In addition, our data reveal that downregulation of c-FLIP promotes the survival of ECM-detached cells and that c-FLIP overexpression in cancer cells restricts the viability of cancer cells grown in anchorage-independent conditions. Taken together, our study reveals an unexpected role for c-FLIP in constraining the viability of cancer cells during ECM-detachment and raises the idea that c-FLIP may have context-dependent pro- and anti-cell death roles during tumorigenesis.

## Introduction

Cellular FLICE (FADD-like IL-1β-converting enzyme)-inhibitory protein (c-FLIP) is a member of the death effector domain (DED) family of proteins along with FADD and pro-caspase 8^1^. While there are multiple c-FLIP variants known to be generated by alternative splicing, each variant has two DED domains and can interact with other DED-containing proteins through DED:DED interactions. These interactions drive the best characterized function of c-FLIP: its capacity to inhibit the activation of receptor-mediated cell death pathways^2^. Mechanistically, this occurs primarily by preventing homodimerization and activation of pro-caspase 8 on the death-inducing signaling complex (DISC). c-FLIP can also function to block necroptosis and promote cell survival by promoting the cleavage of RIPK1 on the ripoptosome^3,4^. As a protein known to inhibit cell death, antagonizing this function of c-FLIP has long been considered a possible strategy to sensitize cancer cells to cell death^5,6^.

Relatedly, the metastasis of cancer cells to distant sites is the primary cause of mortality in cancer patients^7,8^. For cancer cells to grow and effectively metastasize to distant sites, they must overcome several barriers during tumor progression. One such barrier is the lack of integrin-mediated attachment to the extracellular matrix (ECM), which is critical for the survival of a variety of cell types^9^. The term “anoikis” was coined to describe caspase-mediated cell death caused by lack of ECM-attachment^10^ and cancer cells often disable anoikis in order to facilitate survival during metastasis^8^. In addition to anoikis induction, ECM-detachment results in substantial alterations in cellular metabolism that can compromise the viability of ECM-detached cells in an anoikis-independent (or caspase-independent) fashion^11–15^. Oncogenic signaling cascades have been discovered to result in anoikis inhibition and fundamental changes in metabolism that in aggregate function to permit the survival of ECM-detached cancer cells^16–19^.

Here, we report that breast cancer cells benefit from diminished c-FLIP expression, a surprising result given the well-established anti-apoptotic function of c-FLIP. We discovered that c-FLIP is diminished in breast tumors when compared to normal breast tissue and that c-FLIP expression in breast cancer is inversely correlated with the expression of oncogenes. Furthermore, ECM-detachment functions as a signal to induce c-FLIP expression in non-cancerous mammary epithelial cells. Signal transduction emanating from activated oncogenes lowers the ECM-detachment-mediated elevation in c-FLIP expression through a mechanism dependent on PI(3)K signaling. Diminished ECM-detachment-mediated c-FLIP expression enhances the survival of ECM-detached cells in the luminal space of mammary acini, and elevated c-FLIP expression can compromise the viability of breast cancer cells when grown in anchorage-independent conditions. Taken together, our data suggest a non-canonical role for c-FLIP in compromising the viability of ECM-detached breast cancer cells and that downregulation of c-FLIP may facilitate cancer cell survival. As such, our results unveil a possible mechanism to explicate why breast tumors often have diminished c-FLIP levels and why low c-FLIP levels correlate with poor patient outcomes for breast cancer patients.

## Results

### Patient data reveal lower levels of c-FLIP in breast tumor tissue

Given the aforementioned role of c-FLIP in blocking death receptor-mediated cell death, we reasoned that there would be elevated expression of c-FLIP in tumor tissue when compared to normal tissue. Using publicly available data to compare *CFLAR* (which encodes for c-FLIP) expression in tumors compared to normal tissue, we surprisingly observed that *CFLAR* expression was lower in breast cancer (BRCA) when compared to normal counterpart tissue (Fig. 1A). Similar findings were also observed in several other cancer types (Supplemental Fig. 1A). Given these data, we hypothesized that the downregulation in c-FLIP expression in breast tumor tissue may be a consequence of elevated oncogenic signaling. Indeed, an analysis of data derived from The Cancer Genome Atlas (TCGA) revealed an inverse correlation between the expression levels of oncogenes (*HRAS* and *AKT1)* and *CFLAR* in breast (Fig. 1B,C) and lung cancer samples (Supplemental Fig. 1B,C). Taken together, these data suggest that breast cancer cells benefit from lower *CFLAR* expression and may achieve this outcome as a consequence of oncogenic signaling.

**Figure 1.**
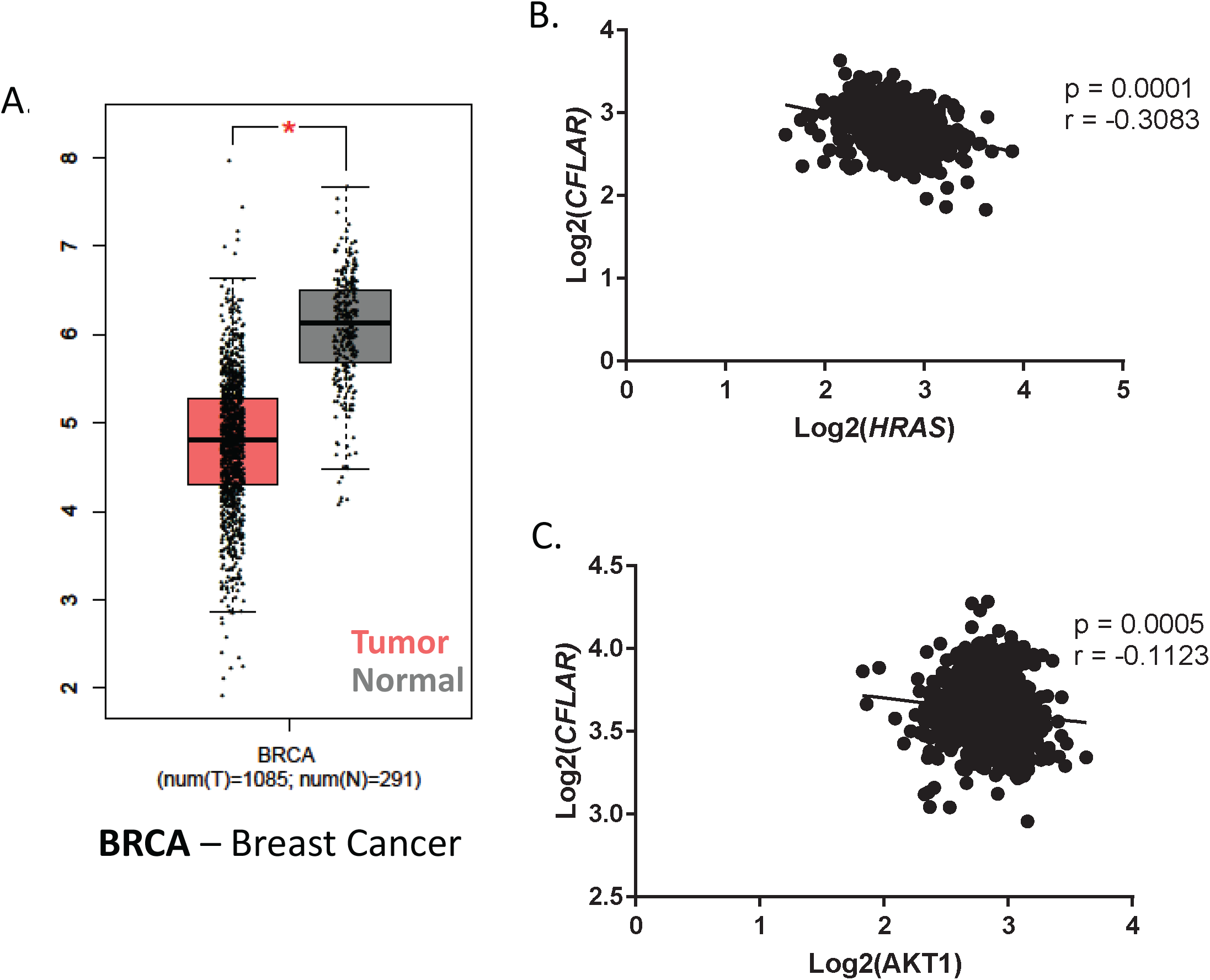
Downregulation of c-FLIP in tumors compared to normal tissue. (A) Comparison of *CFLAR* expression levels in tumor versus normal tissue in breast cancer samples (BRCA). (B and C) Correlative analysis of the expression levels of *CFLAR* with (B) *HRAS* and (C) *AKT1* in breast cancer samples. (A) Data was acquired using online tool known as GEPIA which utilizes TCGA (tumor and normal tissue gene expression) and GTEx (normal tissue gene expression) in order to ascertain relative expression levels of a gene across different tissues (Log2FC cutoff==1, p.value cutoff==0.01); data is presented in Log scale.

### Detachment from ECM causes elevated c-FLIP expression in non-cancerous mammary epithelial cells

Given the surprising evidence that *CFLAR* expression was diminished in breast tumors, we were interested in determining a biological rationale that could lead to such an outcome. To that end, we assessed how c-FLIP was regulated during ECM-detachment. We measured c-FLIP protein levels in established, non-cancerous mammary epithelial cell lines (MCF-10A and HMEC) and found that in both cases, ECM-detachment causes an elevation in c-FLIP protein levels (Fig. 2A). In order to extend our analysis into mammary epithelial cells that more closely resemble cells found in normal mammary tissue, we measured c-FLIP protein levels in KTB34 and KTB37 cells. These immortalized lines were derived from core biopsies of normal breast tissue and, in contrast to MCF-10A or HMEC, display either luminal A (KTB34) or normal-like (KTB37) gene expression patterns^20^. Much like we observed with MCF-10A or HMEC cells, ECM-detachment was a strong signal for c-FLIP induction in KTB34 and KTB37 cells (Fig. 2B). In addition, when we assessed the capacity of ECM-detachment to alter c-FLIP levels in a range of breast cancer lines representing distinct molecular subtypes (Supplemental Fig. 2A), the ability of ECM-detachment to cause c-FLIP upregulation was largely not observed. Furthermore, the elevated c-FLIP levels observed during ECM-detachment are sustained for periods of time (48 hours) known to cause robust caspase activation (Supplemental Figs. 2B,C). Thus, the ECM-detachment-mediated induction in c-FLIP does not result in appreciable anoikis inhibition and may instead have an alternative function during ECM-detachment.

**Figure 2:**
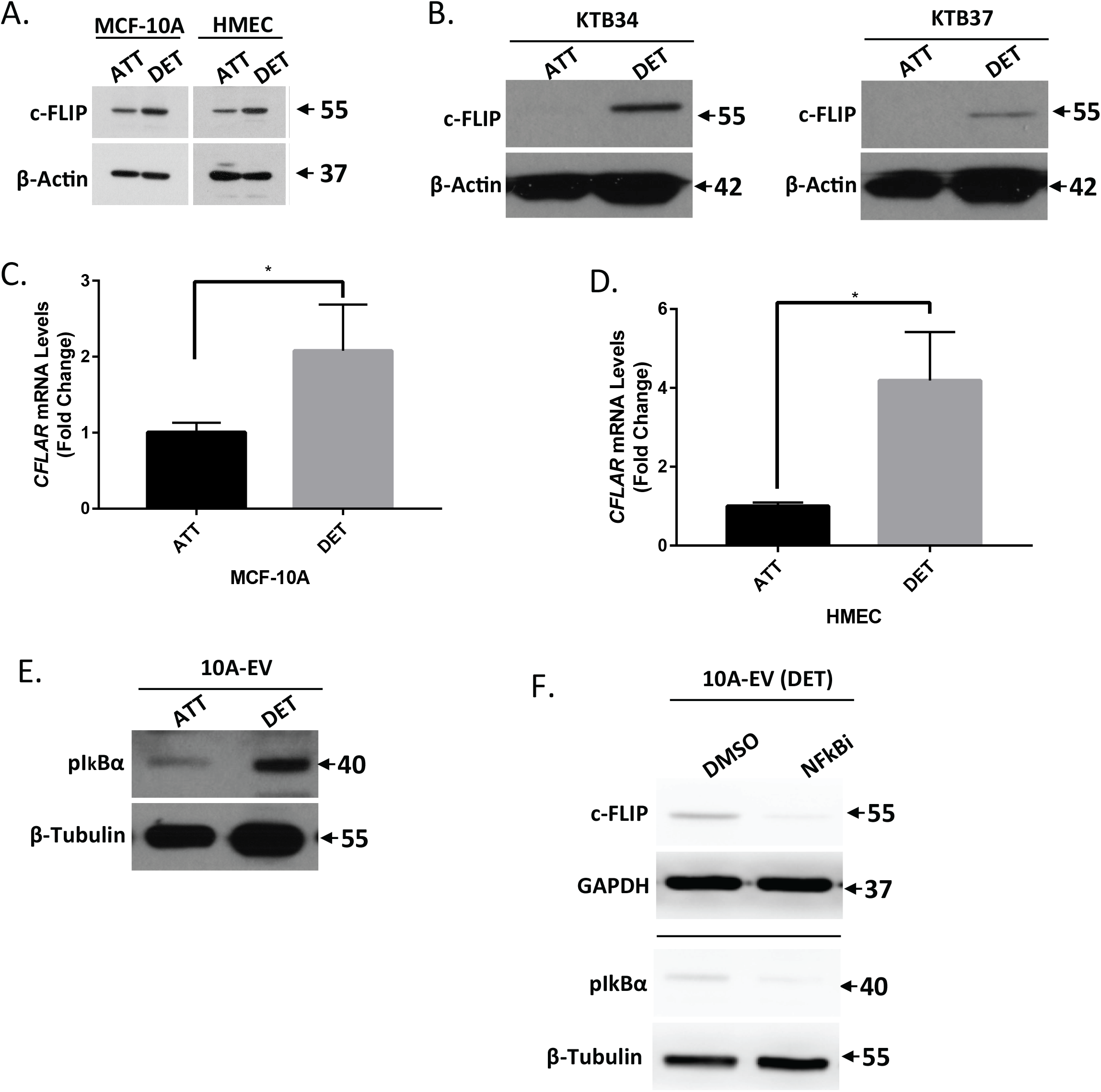
ECM-Detachment triggers c-FLIP expression in non-cancerous epithelial cells. (A) Measurement of c-FLIP protein levels in MCF-10A cells in attachment (ATT) and detachment (DET) conditions using western blot. (B) Measurement of c-FLIP protein levels in KTB34 cells (left) and KTB37 cells (right) after cells were grown in attachment versus detachment conditions for 24h. (C and D) Measurement of c-FLIP transcript levels (*CFLAR)* using RT qPCR in (C) MCF-10A cells and (D) HMEC cells. (E) Measurement of NFkB signaling in 10A-EV cells by blotting for phopho-IκBα in attachment versus detachment conditions. (F) Measurement of the changes in c-FLIP protein levels following inhibition of NFκB activity using 5uM BAY-117082. Graphs show representative data from a minimum of three biological replicates and all western blotting experiments were independently repeated a minimum of three times with similar results. Statistical significance was determined using Student’s two-tailed t-test. Error bars show standard deviation.

Given that levels of c-FLIP protein have been demonstrated to be regulated by numerous mechanisms, we sought to ascertain if c-FLIP levels were elevated during ECM-detachment as a consequence of increased *CFLAR* transcription. Indeed, we observed a robust increase in *CFLAR* transcript when MCF-10A or HMEC cells were grown in ECM-detachment (Fig. 2C,D). Notably, previous studies have revealed that NFκB signaling can function to induce the expression of c-FLIP in various cellular contexts^21^. Thus, we investigated whether ECM-detachment can trigger the activation of NFκB signaling by measuring the phosphorylation of IκBα at Ser32/36, a well-known marker for NFκB activation^22^. Indeed, ECM-detachment resulted in robust phosphorylation of IκBα in MCF-10A cells (Fig. 2E). Furthermore, pharmacological inhibition of NFκB signaling was sufficient to block the ECM-detachment-mediated induction of c-FLIP in cells grown in ECM-detachment (Fig. 2F). Collectively, these data suggest that loss of ECM-attachment results in transcriptional upregulation of c-FLIP due to the activation of NFκB signaling.

### Oncogene overexpression downregulates c-FLIP expression during ECM detachment

Given the data suggesting that *CFLAR* expression is often downregulated in breast tumors (Fig. 1), we reasoned that the introduction of oncogenic signals in non-cancerous mammary epithelial cells may block the ECM-detachment-mediated elevation in c-FLIP. To test this possibility, we engineered MCF-10A cells to express high levels of H-Ras (G12V), ErbB2, or EGFR (Supplemental Fig. 3A). Interestingly, overexpression of each of these oncogenes resulted in the downregulation of c-FLIP protein (Fig. 3A) and *CFLAR* expression (Fig. 3B) in ECM-detached cells. Given that each of these oncogenes can promote activation of the PI(3)K/Akt pathway^23^, we assessed whether activation of PI(3)K or Akt is sufficient to inhibit the ECM-detachment-mediated elevation in c-FLIP. Indeed, we observed that expression of constitutively active PI(3)K (P110α^E545K^) or Akt (myristoylated-Akt) resulted in reduced c-FLIP levels in ECM-detachment (Fig. 3C, Supplemental Fig. 3B). Next, we assessed whether activation of PI(3)K-Akt is necessary to limit c-FLIP levels in ECM-detached cells. Upon treatment with a selective inhibitor of PI(3)K, c-FLIP protein levels were restored in ECM-detached MCF-10A cells expressing H-Ras (G12V), ErbB2, or EGFR (Fig. 3D). Similarly, treatment with the PI(3)K inhibitor restores *CFLAR* expression in MCF-10A cells engineered to have activated oncogenic signaling (Fig. 3E). However, this capacity to elevate *CFLAR* expression does not extend to control MCF-10A cells that have not been transduced with activated oncogenes (Fig. 3E).

**Figure 3.**
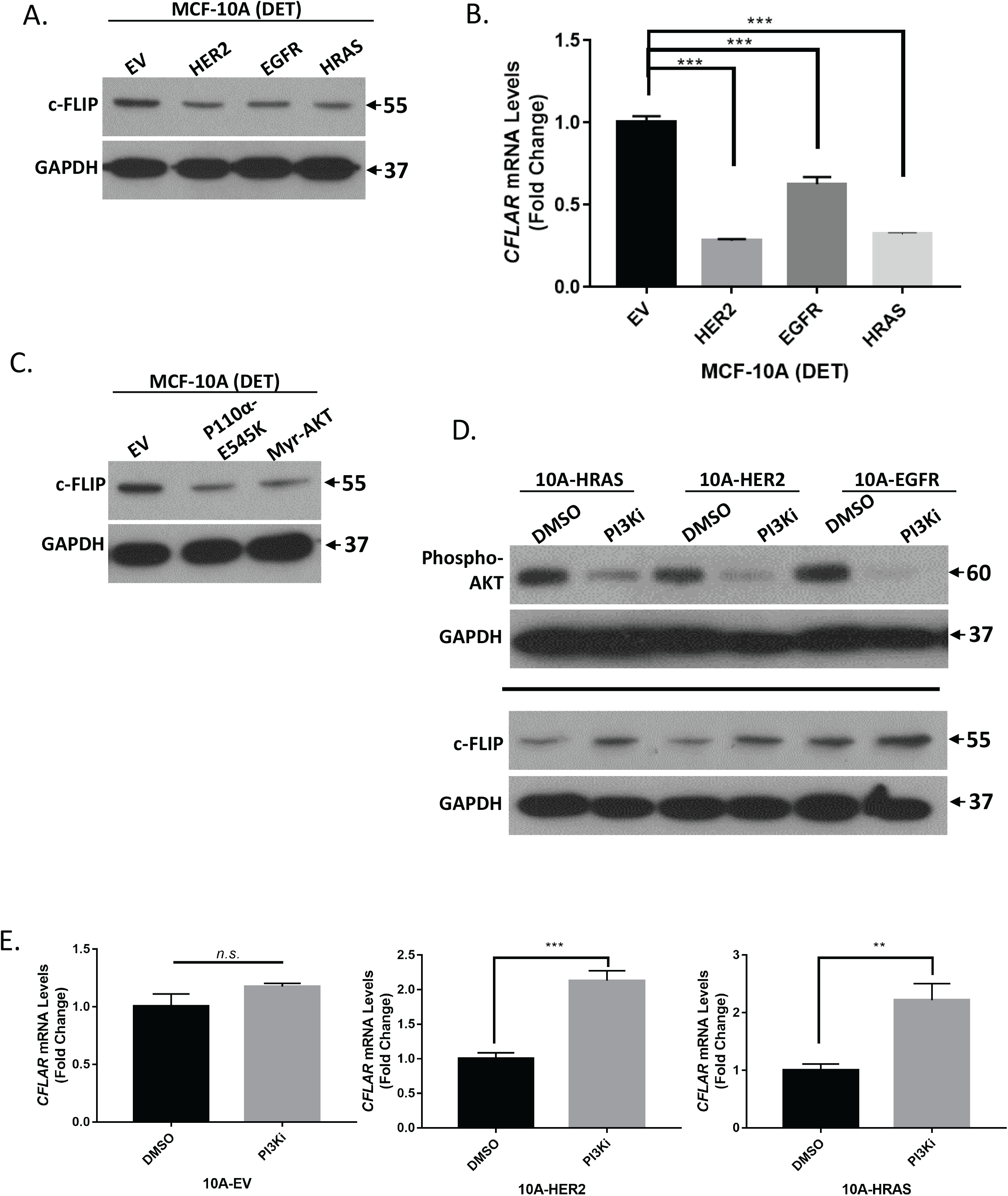
Regulation of c-FLIP by PI(3)K/Akt signaling during ECM-detachment. (A) Measurement of c-FLIP protein levels in MCF-10A cells overexpressing oncogenes (HRAS^G12V^, HER2, EGFR) as compared to empty vector (EV) control. (B) RT-qPCR measurement of *CFLAR* in MCF-10A cells overexpressing oncogenes versus EV control. (C) Measurement of c-FLIP protein levels in MCF-10A cells overexpressiing oncogenes (p110α^E545K^ and myr-AKT) as compared to EV control. (D) Measurement of c-FLIP levels after treatment of oncogene overexpressing MCF-10A cells with small molecule inhibitor of PI3Kα, BYL719 (10uM). (E) Measurement of *CFLAR* levels in oncogene overexpressing MCF-10A cell treated with BYL719. Graphs show representative data from a minimum of three biological replicates and all western blotting experiments were independently repeated a minimum of three times with similar results. Statistical significance was determined using Student’s two-tailed t-test. Error bars show standard deviation. All panels show cells grown in ECM-detached conditions.

### c-Myc is a downstream effector of PI(3)K/Akt that is associated with the downregulation of c-FLIP expression

Given that ECM-detachment-mediated elevation in c-FLIP is abrogated by PI(3)K/Akt signaling, we sought to elucidate the relationship between Akt and c-FLIP. To do so, we first utilized a bioinformatic approach to predict relationships between *AKT1* and *CFLAR* gene expression^24^. This analysis revealed that c-Myc could be an intermediary linking PI(3)K/Akt activation and downregulation of *CFLAR* expression (Fig. 4A). Previous studies have discovered that c-Myc can function as a transcriptional repressor^25,26^ and that in certain cellular contexts, c-Myc can directly repress *CFLAR* expression^27^. Interestingly, we found that that activation of oncogenic signaling (via expression of H-Ras (G12V) or HER2) led to increased c-Myc levels in ECM-detached cells (Fig. 4B). Furthermore, this upregulation is likely due to the stabilization of c-Myc protein, as we do not detect differences in expression of *MYC* mRNA upon activation of oncogenic signaling (Fig. 4C).

**Figure 4.**
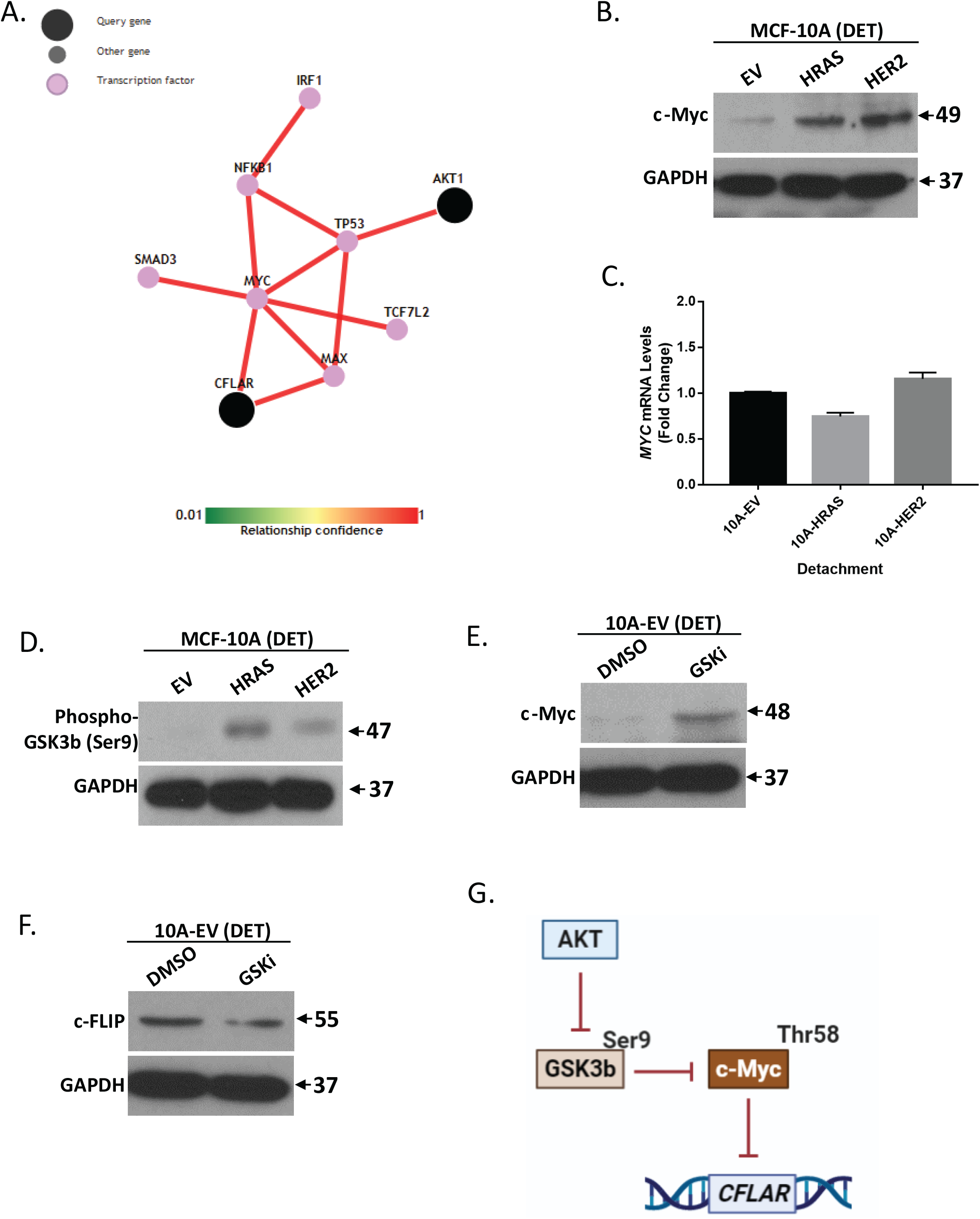
Transcriptional Regulation of c-FLIP during ECM-Detachment. (A) *In-silico* analysis of the transcriptional relationship between *CFLAR* and *AKT1* using a pathway prediction tool (http://pathwaynet.princeton.edu/). (B) Measurement of c-Myc levels in oncogene overexpressing MCF-10A cells as compared to EV control. (C) Measurement of c-Myc transcript levels using RT-qPCR in oncogene overexpressing MCF-10A cells as compared to EV control. (D) Measurement of GSK-3β activity by blotting for its phosphorylation at serine 9. (E) Inhibition of GSK-3β activity using 25uM TDZD-8 (GSK-3β inhibitor) in 10A-EV cells. (F) Determination of c-FLIP protein levels in 10A-EV cells after the inhibition of GSK-3β activity using 25uM TDZD-8. (G) Schematic representation of transcriptional regulation of c-FLIP by Akt. Graphs show representative data from a minimum of three biological replicates and all western blotting experiments were independently repeated a minimum of three times with similar results. Cells were grown in ECM-detached conditions in all panels. Statistical significance was determined using Student’s two-tailed t-test. Error bars show standard deviation.

One prominent and well-described regulator of the stability of c-Myc protein is GSK-3β, which phosphorylates c-Myc at threonine 58 (Thr58). This phosphorylation event is known to trigger the recruitment of E3 ubiquitin ligases and to cause proteasomal degradation of the c-Myc protein^28^. Furthermore, GSK-3β activity is negatively regulated by Akt-mediated phosphorylation at serine 9 (Ser9)^29^. As such, we reasoned that the PI(3)K-mediated downregulation of c-FLIP during ECM-detachment may be a consequence of GSK-3β inhibition and thus stabilization of c-Myc protein. In support of this possibility, we observed increased phosphorylation of GSK-3β at Ser9 in cells engineered to activate oncogenic signaling (Fig. 4D). In addition, treatment of ECM-detached cells with TDZD-8, a GSK-3β inhibitor, led to elevated c-Myc protein levels (Fig. 4E) and diminished abundance of c-FLIP (Fig. 4F). Taken together, these data support a model in which PI(3)K/Akt blocks GSK-3β activity, stabilizes c-Myc protein, and represses transcription of *CFLAR* during ECM-detachment (see model in Fig. 4G).

### c-FLIP antagonizes mammary cell survival during ECM-detachment

Given the aforementioned data demonstrating that oncogenic signaling can downregulate c-FLIP levels during ECM-detachment, we were interested in understanding if loss of c-FLIP would confer a benefit to cells grown in ECM-detachment. As such, we utilized lentiviral delivery of shRNA to engineer MCF-10A cells to be deficient in c-FLIP (Fig. 5A). When grown in Matrigel, MCF-10A cells will form 3-dimensional acinar structures that closely model mammary morphogenesis^30^. In addition, the hollowing of mammary acini is well-known to be controlled by cell death programs activated in centrally located cells that lack attachment to ECM^11,14,16,31,32^. Intriguingly, shRNA-mediated reduction of c-FLIP caused a significant increase in the number of mammary acini that are scored as “mostly filled” or “filled” (Fig. 5B). Thus, we conclude that downregulation of c-FLIP can promote the survival of ECM-detached cells in the luminal space of mammary acini.

**Figure 5.**
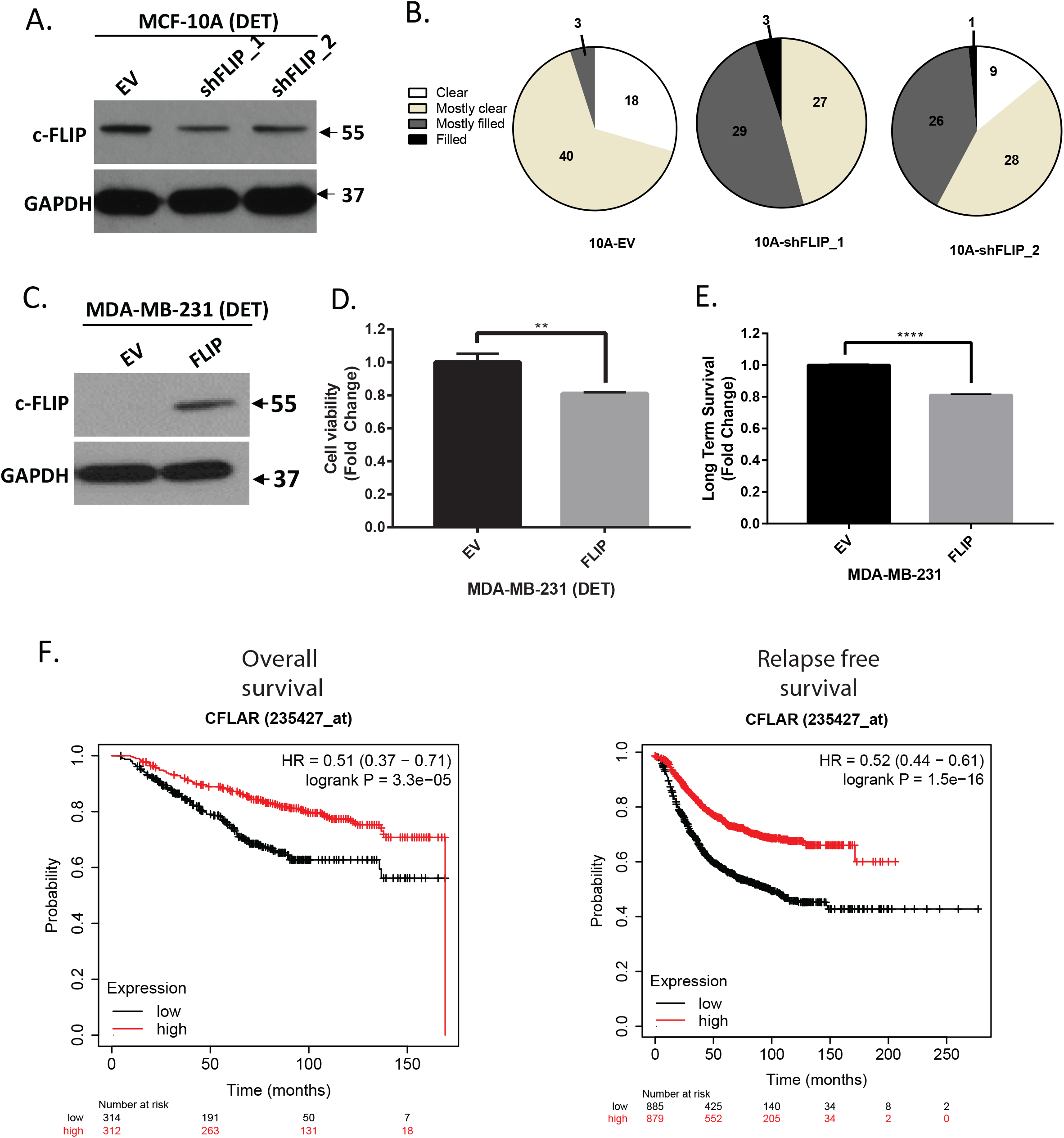
c-FLIP impacts the survival of ECM-detached cells. (A) MCF-10A cells were stably transfected with shRNA targeting c-FLIP and efficacy was verified with western blot. (B) 10A-EV, 10A-shFLIP1 and 10A-shFLIP3 cells were grown in 3D Matrigel Assay and were allowed to form acinar structures for 18 days. Structures were stained and luminal clearance was scored as described. 10A-EV *n* = 61, 10A-shFLIP1 *n* = 59, 10A-shFLIP3 *n* = 64. (C) c-FLIP was overexpressed in MDA-MB-231 cells and verified by western blot. (D) Measurement of cell viability MDA-MB-231 cells during ECM-detachment using CellTiter-Glo^®^ Luminescent Cell Viability Assay after 48 hours. (E) Long term survival assay of MDA-MB-231 cells tested as described and stained with crystal violet assay. Absorbance of extracted crystal violet at 590 nM is shown. (F) Correlation of c-FLIP expression to overall survival and relapse-free survival of patients with breast cancer. c-FLIP expression is differentiated as low (black) vs high (red) expression against the median expression. Overall survival was analyzed using KMPlotter (www.kmplot.com). Graphs show representative data from a minimum of three biological replicates and all western blotting experiments were independently repeated a minimum of three times with similar results. Statistical significance was determined using Student’s two-tailed t-test. Error bars show standard deviation.

Given that our data suggest that loss of c-FLIP can promote the survival of non-cancerous mammary epithelial cells grown in ECM-detached conditions, we reasoned that elevation of c-FLIP levels in invasive breast cancer cells may compromise their survival. To test this possibility, we overexpressed c-FLIP in MDA-MB-231 cells, a highly aggressive, triple negative breast cancer line (Fig 5C). Indeed, c-FLIP compromised the viability of these cells when grown in ECM-detached conditions (Fig. 5D). Similarly, when cells were grown in ECM-detachment and then re-plated at low density in ECM-attached conditions, c-FLIP expression blocked the capacity of these cells to form colonies (Fig. 5E). Given the data that lower levels of c-FLIP may promote the survival of ECM-detached cancer cells, we hypothesized that low levels of c-FLIP may be associated with poor clinical outcomes owing to tumor cell dissemination. Indeed, analysis of data derived from breast cancer patients revealed that lower levels of *CFLAR* expression was linked to poor patient outcomes as measured by diminished overall survival and relapse-free survival (Fig. 5F).

## Discussion

Our findings describe an unexpected role for c-FLIP during ECM-detachment that may account for the observed downregulation of c-FLIP in breast cancers. Furthermore, our results demonstrate that the ECM-detachment-mediated elevation in c-FLIP expression is counteracted by oncogenic signaling through activation of the PI(3)K/Akt pathway. Oncogenic activation of PI(3)K/Akt is associated with inhibition of GSK-3β and an elevation in c-Myc-mediated downregulation of *CFLAR* expression. In support of these data, we found that diminished *CFLAR* expression (which functions to facilitate the survival of ECM-detached cells) is correlated with poor clinical outcomes in breast cancer patients. As such, our studies reveal that restoration of this novel c-FLIP activity may be an attractive chemotherapeutic strategy to eliminate ECM-detached cancer cells prior to (or during) metastatic dissemination.

Our findings raise interesting questions regarding dichotomous and context-dependent roles for c-FLIP during the course of tumorigenesis. More specifically, there appears to be substantial evidence of downregulation of c-FLIP in breast tumors where low c-FLIP is also associated with oncogenic signaling and with poor patient outcomes. The ability of c-FLIP to antagonize the viability of ECM-detached cancer cells may be related to the observed changes in c-FLIP in tumors derived from breast cancer patients. Additionally, we do have data (see Supplemental Fig. 1) that reveal some other cancers also have diminished c-FLIP levels compared to normal tissue. Future studies aimed at broadening an assessment of c-FLIP in tumors of distinct origins, and the relationship between oncogenic signaling and c-FLIP in other types of cancer cells, will be important for better understanding the implications of these findings.

Furthermore, any efforts to therapeutically restore c-FLIP activity to restrict tumor progression would have to contend with the context-dependent nature of c-FLIP activity. The capacity of c-FLIP to block cell death by apoptosis has been thoroughly described and thus, it is important to have a better understanding of the molecular circumstances that underlie when c-FLIP can be pro-tumorigenic compared to when c-FLIP can be anti-tumorigenic (as may be the case during ECM-detachment).

In addition, the contrast between c-FLIP function in ECM-detachment and in other contexts is stark. The well-characterized anti-apoptotic roles of c-FLIP are, on the surface, difficult to reconcile with the apparently pro-death function of c-FLIP described in ECM-detachment. However, there are indeed other reported instances where c-FLIP has been demonstrated to have multiple roles with regards to cell death regulation. For example, c-FLIP null mice do not survive beyond day 10.5 of embryogenesis due to defects in heart development^33^. This phenotype is similar to that observed of FADD −/− or caspase 8 −/− mice, suggesting that c-FLIP function can align with the function of pro-cell death proteins during embryonic development. Our data raise interesting future questions about the precise mechanism(s) employed by c-FLIP to compromise the viability of ECM-detached cells. Interestingly, c-FLIP levels can be regulated by glutamine starvation^34^, and c-FLIP can promote SGLT1-mediated glucose uptake in hepatocellular carcinoma cells^35^. Given these data and the fact that ECM-detachment is a profound signal to alter nutrient uptake and utilization, it is possible that the function of c-FLIP during ECM-detachment involves alterations in cellular metabolism. Future studies aimed at better understanding the nexus between ECM-detachment, c-FLIP, and viability will be important in order to better grasp the multi-faceted role played by c-FLIP in cancer pathogenesis.

## Methods

### Cell culture

MCF-10A cells (ATCC, Manassas, VA, USA) and derivatives were cultured in Dulbecco’s Modified Eagle Medium/F12 (Gibco, Waltham, MA, USA) supplemented with 5% horse serum (Invitrogen), 20ng/mL epidermal growth factor (EGF), insulin (10μg/mL), hydrocortisone (500μg/mL), cholera toxin (100ng/mL), and 1% penicillin/streptomycin. HMEC-Human Mammary Epithelial Cells (Lonza, Basel, Switzerland) and derivatives were cultured in Mammary Epithelium Basal Medium (MEBM) (Lonza) plus MEGM BulletKit and 1% penicillin/streptomycin. MDA-MB-231 cells (ATCC) and derivatives were cultured in Dulbecco’s Modified Eagle’s Medium with 10% fetal bovine serum (Invitrogen) and 1% penicillin/streptomycin. The small molecule inhibitors used are as follows: BYL719 (APExBIO, A8346), TDZD-8 (Enzo life sciences, ALX-270-354-M005) and BAY-117082 (Sigma Aldrich, B5556). Cells were grown for 24 hours unless stated otherwise.

### Immunoblotting

ECM-detached cells were harvested, washed once with cold PBS, and lysed in 1% Nonidet P-40 supplemented with protease inhibitors leupeptin (5μg/mL), aprotinin (1μg/mL), and PMSF (1mM) and the Halt Phosphatase Inhibitor Mixture (Thermo Scientific, Waltham, MA, USA). Lysates were collected after spinning for 30 minutes at 4°C at 14,000 rpm and normalized by BCA Assay (Pierce Biotechnology, Waltham, MA, USA). Normalized lysates underwent SDS-PAGE and transfer/blotting was performed as previously described (Davison et al., 2013). The following antibodies were used for western blotting: FLIP (Cell Signaling Technology, #56343), phospho-Akt (Ser473) (Cell Signaling Technology, #4060), GAPDH (Cell Signaling Technologies, #5174), β-tubulin (Cell Signaling Technology, #2146), β-Actin (Sigma-Aldrich, #A1978), phospho-IκBα (Ser32/36) (Cell Signaling Technology, 9246s), phospho-GSK-3β (Ser9) (Cell Signaling Technology, 5558s), and c-myc (sigma M-5546).

### RNA isolation and quantitative real-time PCR

Total RNA was isolated with RNeasy Mini Kit (Qiagen, Germantown, MD, USA). RNA (1 μg) was reverse transcribed into cDNA using iScript Reverse Transcription Supermix Kit (Bio-Rad, Hercules, CA, USA). The relative levels of gene transcripts compared to the control gene 18S were determined by quantitative real-time PCR using SYBER Green PCR Supermix (Bio-Rad) and specific primers on a 7,500 Fast Transient transfection Real Time PCR System (Applied Biosystems, Life Technologies). Amplification was performed at 95°C for 12 min, followed by 40 cycles of 15 s at 95°C, and 1 min at 60°C. The fold change in gene expression was calculated as: Fold change = 2^-ddCT^. The primers used are as follows: *18S* primers (F – GGCGCCCCCTCGATGCTCTTAG; R - GCTCGGGCCTGCTTTGAACACTCT), *CFLAR* primers (F - GTGGAGACCCACCTGCTCA; R - GGACACATCAGATTTATCCAAATCC), *MYC* primers (F – AGGGTCAAGTTGGACAGTGTCA; R – TGGTCGATTTTCGGTTGTTG), *IL6* primers (F - ACATCCTCGACGGCATCTCA; R - TCACCAGGCAAGTCTCCTCA).

### Lentiviral delivery of shRNA and generation of stable cell lines

MISSION short hairpin RNA (shRNA) constructs against c-FLIP (TRCN0000007228 and TRCN0000007230) in the puromycin-resistant pLKO.4 vector along with an empty vector control were purchased from Sigma-Aldrich. HEK293T cells were transfected with 0.5μg target DNA along with the packaging vectors pCMV-D8.9 (0.5μg) and pCMV-VSV-G (60ng) using Lipofectamine 2000 and PLUS reagent (Life Technologies). Virus was collected 24- and 48-hours post-transfection and filtered through a 0.45μm filter (EMD Millipore) and used for transduction of MCF-10A cells in the presence of polybrene (8μg/mL). Stable populations of MCF-10A cells were selected using puromycin (2μg/mL) (Invivogen, San Diego, CA, USA).

### Retroviral mediated generation of stable cell lines

The pBABE-Puro-based retroviral vectors encoding constitutively active HRAS^G12V^, EGFR, ERBB2, constitutively active P110α^E545K^, and myristoylated-AKT were used to generate stable cell lines. The pBABE-Puro-based retroviral vectors encoding FLIP was used to generate stable cell lines. HEK293T cells were transfected with target DNA (0.75μg) along with the packaging vector pCLAmpho (0.75μg) with Lipofectamine 2000 (Life Technologies). Virus was collected at 48- and 72-hours post-transfection, filtered through a 0.45μm filter (EMD Millipore), and used for transduction of MCF-10A and MDA-MB-231 cells in the presence of polybrene (8μg/mL). Stable populations of puromycin-resistant cells were obtained using puromycin (2μg/mL) (Invivogen, San Diego, CA, USA).

### Caspase assay

Cells were plated at a density of 5,000 cells per well on 96-well plates. Caspase activation was measured using the Caspase Glo 3/7 Assay Kit (Promega, Madison, WI, USA) according to manufacturer’s instructions.

### Cell viability assay

Cells were plated at a density of 5,000 cells per well on 96-well plates. Caspase activation was measured using the Cell Titer Glo Assay Kit (Promega, Madison, WI, USA) according to manufacturer’s instructions.

### Crystal Violet assay

MDA-MB-231 Cells were plated at 100,000 cells per well in poly-HEMA coated 6-well plates for 96 hours. After 96 hours, cells were washed and were transferred to adherent 6-well plates for 24 hours. Cells were then washed with 1x PBS and then were fixed and stained with 750 μl crystal violet solution (0.5%) for 10 min. Cells were washed with deionized water. After imaging plates, cells were destained by adding 1 ml of 10% acetic acid per well and plates were rocked for 20 min at room temperature. Samples from each well were transferred to a 96-well plate, and absorbance at 590 nM was read using a Spectramax M5 plate reader (Molecular Devices). Statistical analysis of absorbance at 590 nM was performed using two-way ANOVA.

### TCGA data analysis

#### Comparison of c-FLIP levels based on clinical attributes

The dataset used was “Breast Invasive Carcinoma (TCGA, Firehose Legacy)”. mRNA expression data of breast cancer datasets with specific clinical attributes (ER status and PR status) was acquired. Statistical analysis on the comparison of CFLAR expression levels to receptor status was performed using two-way ANOVA.

#### Correlation of CFLAR expression to oncogenes

The dataset used were “Breast Invasive Carcinoma (TCGA, Firehose Legacy)” and “Lung Adenocarcinoma (TCGA, Firehose Legacy)”. Co-expression data of *CFLAR* and selected genes (HRAS, AKT1, ERBB2) was acquired from the database. Statistical analysis on the comparison of CFLAR expression levels to receptor status was performed using two-way ANOVA.

## Acknowledgements

We thank Veronica Schafer and all current and past Schafer laboratory members for helpful comments, experimental assistance, and/or valuable discussion. We also thank the Siyuan Zhang lab (Notre Dame) and Xin Lu lab (Notre Dame) for technical and conceptual feedback. We thank the Sandra Van Schaeybroeck lab (Queen’s University Belfast, Belfast, UK) for sharing the c-FLIP overexpression plasmid and Harikrishna Nakshatri (Indiana University School of Medicine) for sharing the KTB cell lines. The graphic in Figure 4G was created with Biorender.com. This work was supported by a Career Catalyst Research Grant (CCR14302768) from Susan G. Komen (to Z.T.S.), a Lee National Denim Day Research Scholar Grant (RSG-14-145-01) from the American Cancer Society (to Z.T.S.), the Coleman Foundation, the College of Science at the University of Notre Dame, and the Malanga Family Excellence Fund for Cancer Research.

## Author Contributions

M.A.T., J.H., K.M., I.M.M. and M.N. conducted experiments. M.A.T., J.H., K.M., I.M.M., M.N., and Z.T.S. analyzed data and interpreted results. M.A.T. and Z.T.S. wrote the manuscript with feedback from all other authors. Z.T.S. was responsible for conception/design of the project and overall study supervision.

## Additional information

The authors declare no competing interests.

## Notes

### Competing Interest Statement

The authors have declared no competing interest.

