## Supplementary Figures 1-3 for "Oncogenic signaling inhibits c-FLIP expression and promotes cancer cell survival during ECM-detachment"

A.

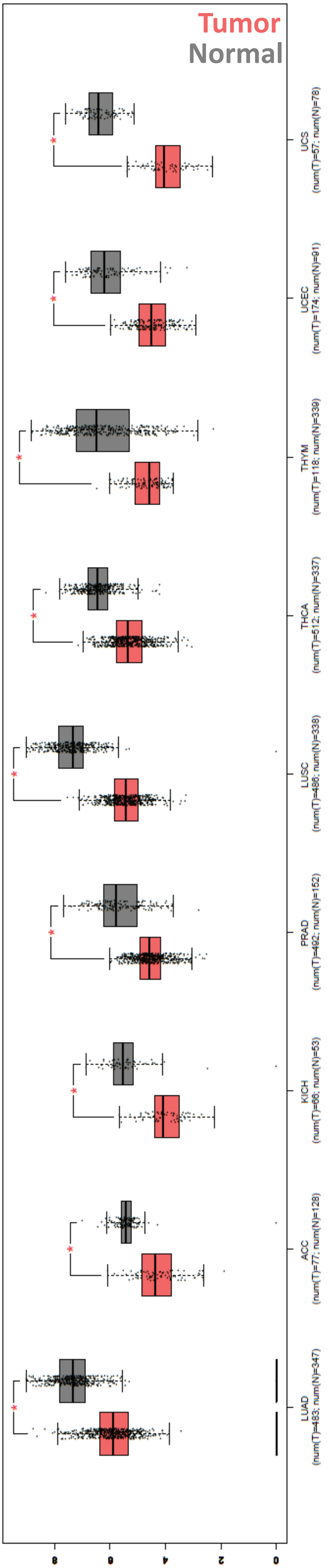


**LUAD** Lung Adenocarcinoma
**ACC –** Adenocortical carcinoma
**KICH –** Kydney Chromophobe
**PRAD** – Prostate Adenocarcinoma
**LUSC** – Lung Squamous Cell Carcinoma
**THCA** – Thyroid carcinoma
**THYM** – Thymoma
**UCEC** – Uterine Corpus Endometrial Carcinoma
**UCS –** Uterine Carcinosarcoma

B.

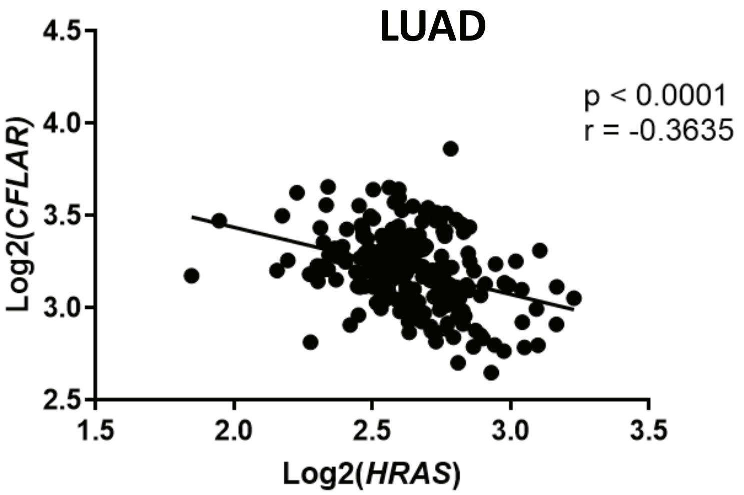


C.

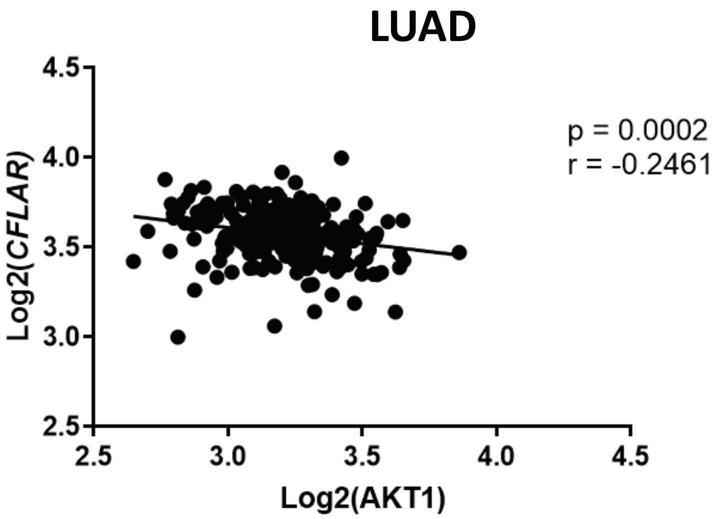


**Supplemental Figure 1**

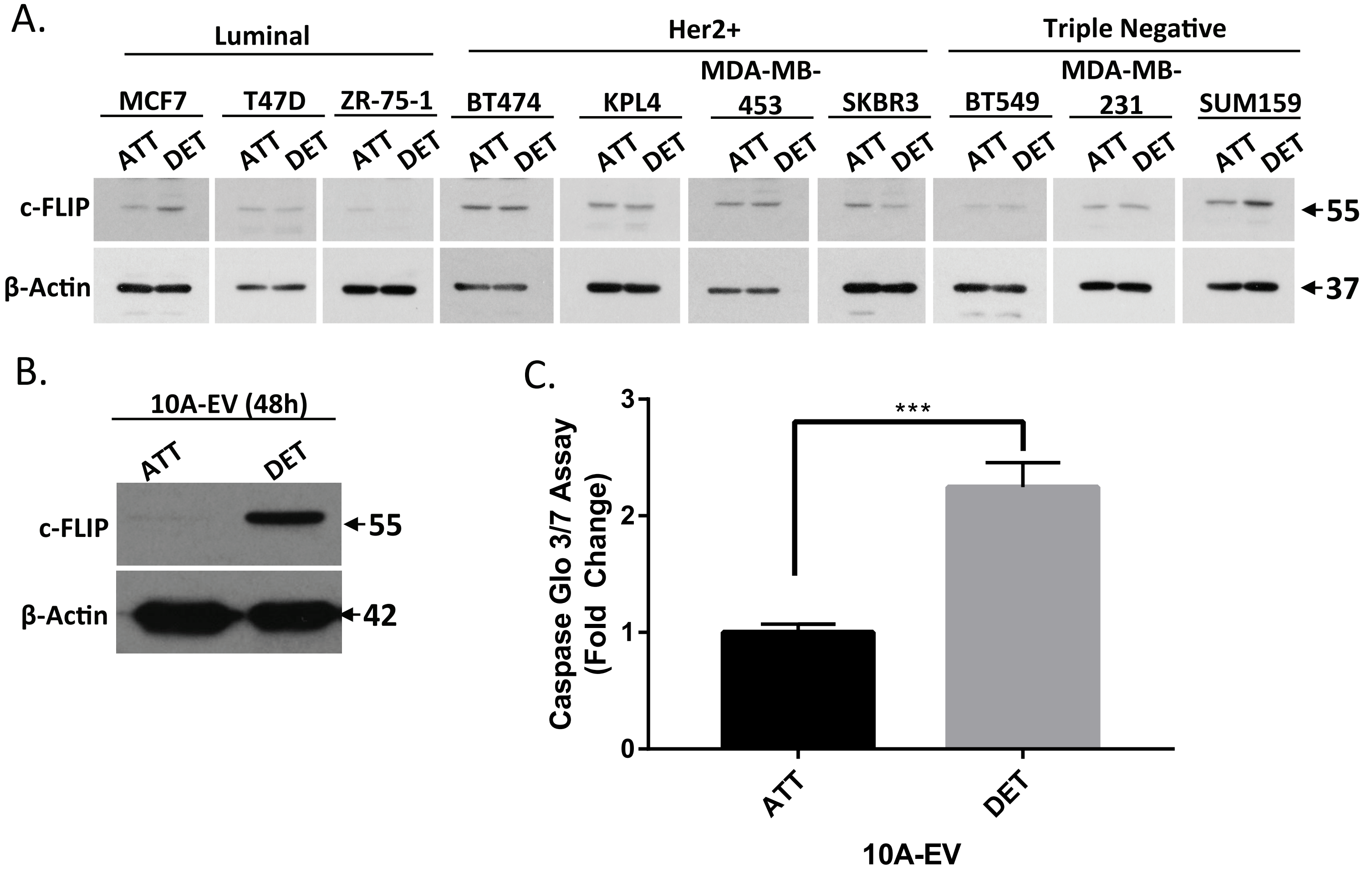


### Supplemental Figure 2

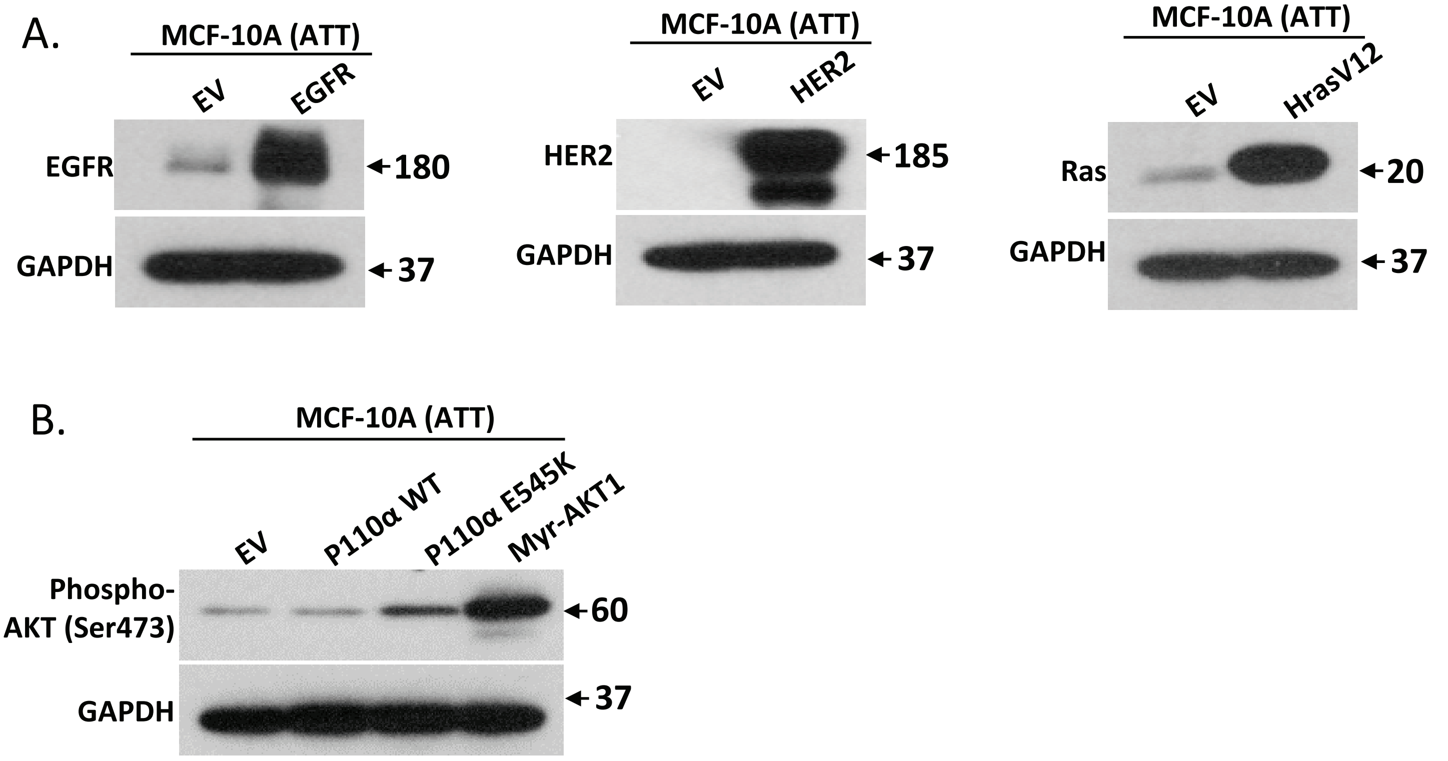


### Supplemental Figure 3

#### Supplemental Figure Legends

**Supplemental Figure 1. Downregulation of c-FLIP is observed in different cancer types**

(A) Comparison of *CFLAR* expression levels in tumor versus normal tissue in different cancer types; (B and C) Correlative analysis of the expression levels of *CFLAR* with (B) *HRAS* and (C) *AKT1* in lung cancer samples.

**Supplemental Figure 2. Expression of c-FLIP during ECM-detachment is independent of caspase 3 activation**

(A) Comparison of c-FLIP protein levels in attachment versus detachment conditions in cell lines representing different subtypes of breast cancer. (B) Measurement of c-FLIP levels in 10A-EV cells after being grown for 48h in ECM-detached condition. (C) Measurement of Caspase 3/7 activity in 10A-EV cells after being grown for 48h in ECM-detached condition. Statistical significance was determined using Student's two-tailed t-test. Error bars show standard deviation. Western blots and biochemical assays show representative data from three biological replicates. Statistical significance was determined using Student's two-tailed t-test. Error bars show standard deviation.

**Supplemental Figure 3. Regulation of c-FLIP is primarily driven by PI(3)K/Akt signaling**

(A) Verification of EGFR (left), HER2 (middle), H-Ras G12V (right) overexpression in MCF-10A cells. (B) Confirmation of PI(3)K activation in indicated cell lines via immunoblotting of phopsho-Akt (Ser473). Western blots show representative data from three biological replicates.
